# Persistent increases of PKMζ in sensorimotor cortex maintain procedural long-term memory storage

**DOI:** 10.1101/193508

**Authors:** Peng P. Gao, Jeffrey H. Goodman, Todd C. Sacktor, Joseph T. Francis

## Abstract

Procedural memories, such as for riding a bicycle, can be maintained without practice for long periods of time and are thought to be supported by the persistent reorganization of sensorimotor cortices (S1/M1). Whereas enhanced synaptic strength and structural changes accompany the learning of motor tasks, the persistent molecular modifications that store long-term procedural memories within specific layers of sensorimotor cortex have not been identified. The persistent increase in the autonomously active, atypical PKC isoform, PKMζ, is a putative molecular mechanism for maintaining enhanced synaptic strength during long-term potentiation (LTP) and several forms of long-term memory. Here we examine whether persistent increases in PKMζ store long-term memory for a reaching task in rat sensorimotor cortex that could reveal the sites of procedural memory storage. Perturbing PKMζ synthesis with PKMζ -antisense oligodeoxynucleotides or blocking atypical PKC activity with zeta inhibitory peptide (ZIP) in S1/M1 disrupts and erases the maintenance of long-term motor memories. Only memories that are maintained without daily reinforcement are affected, indicating atypical PKCs (via ZIP) and PKMζ specifically (via antisense) stores consolidated long-term procedural memories. Analysis of changes in the amount of PKMζ in S1/M1 reveals PKMζ increases in layers II/III and V of both S1 and M1 cortices as performance improves to an asymptote during training. After storage for 1 month without reinforcement, the increase in M1 layer V but not other layers persists without decrement. Thus, the sustained increases in PKMζ reveal that the persistent molecular changes storing long-term procedural memory are localized to the descending output layer of primary motor cortex.

## Introduction

Motor learning is characterized by a slow improvement of the smoothness and accuracy of skilled movements, which, once established, are maintained for long periods of time without further practice [1]. A skilled reaching task in which rodents are trained to reach with their preferred forelimb through a small slot to grasp food pellets has been widely used to study the neural substrate underlying motor learning [2-11]. Performance gains and the maintenance of proficiency on this task depend on the integrity of the sensorimotor cortex [5,12-14]. Plastic changes in sensorimotor cortex, including synaptic strength modification and structural remodeling, have been correlated with different phases of the learning process [4,8-11,15,16]. Rioult-Pedotti and colleagues, for example, found the synaptic efficacy of horizontal connections in primary motor cortex (M1) layer II/III increased significantly, after 5 days of training, on the contralateral hemisphere to the preferred forelimb [11], indicating an LTP-like modification of synaptic transmission [8,9]. Both spine formation and elimination were seen right after the first motor learning session in mice [15] and sustained for up to 20 days with continued daily training in rodents [15,17]. However, the molecular mechanisms that store motor memories in sensorimotor cortex remain unknown.

The persistent increase in the autonomously active, atypical PKC (aPKC) isoform PKMζ is both necessary and sufficient for maintaining LTP [18-20]. PKMζ activity retains increased amounts of α-amino-3-hydroxy-5-methyl-4-isoxazolepropionic acid-type glutamate receptors (AMPARs) in postsynaptic sites to maintain synaptic potentiation [21,22]. PKMζ also contributes to maintaining the structural modifications of dendritic spines and synapses, changes which have been extensively observed in sensorimotor cortex after sensorimotor learning [23,24]. Inhibition of persistent atypical PKC activity in specific brain structures by zeta inhibitory peptide (ZIP) disrupts the maintenance of various types of memory, including hippocampus-dependent spatial memory [25], basolateral amygdala-dependent fear memories [21,26-29], dorsal lateral striatum-dependent habit memory [30] and insular cortex-dependent long-term associative conditioned taste aversion memory [31], as well as sensorimotor cortex-dependent motor learning in the reaching task [32]. However, ZIP affects PKC isoforms in addition to PKMζ, particularly the other atypical PKC, PKC_ι/λ_, which can compensate for PKMζ in PKMζ-knockout mice [33]. Therefore, we started by investigating the role of PKMζ in motor learning with PKMζ -antisense oligodeoxynucleotides directed against the translational start site of the PKMζ mRNA, which is specifically disrupts PKMζ synthesis and not PKC_ι/λ_ synthesis [33].

## Results

### Learning phases of a skilled reaching task

We used a skilled reaching task to study the role of PKMζ in sensorimotor learning and long-term memory maintenance. Rats were trained to reach with their preferred forelimbs through a small slot to grasp food pellets. Repeated training sessions were required to obtain good performance, which provided an extended time window to examine in detail any changes in sensorimotor cortex. The success rate, defined as % successful reaches / total reaching attempts, in a 30-minute training session, was used to evaluate task performance on a daily basis. A successful reach includes: 1) lift and advance the preferred forelimb through the slot; 2) pronate and grasp food pellet; 3) retrieve without dropping the pellet [34].

Consistent with the model described by Monfils and Teskey [8], we observed that the acquisition and maintenance of skilled motor memory can be divided into four phases (Figure S 1): 1) a skill acquisition phase (Days 1-4), when success rate is comparatively low (<30%); 2) a performance improvement phase (Days 5-9), when the success rate increases rapidly until plateauing (~70%); 3) the proficiency maintenance phase, when performance is stable with continual training; and 4) a long-term memory storage phase, when performance is stable without training. Because ZIP appears to specifically disrupt the long-term sensorimotor memory storage phase [32], we first focused on the effects of PKMζ-antisense on the acquisition and subsequent maintenance of sensorimotor learning and memories.

### PKMζ-antisense slows the performance improvement phase of sensorimotor learning

To determine the necessity of PKMζ in sensorimotor cortical networks during sensorimotor learning and memory maintenance, we utilized antisense oligodeoxynucleotides against the translation start site of PKMζ mRNA, which effectively blocks PKMζ synthesis and has no effect on LTP or memory formation in the absence of its specific target PKMζ mRNA [33,35]. We injected PKMζ-antisense bilaterally in S1/M1 30 minutes before daily training and examined its effect on sensorimotor learning (Figure 1 and Figure S 2). The antisense group had a significantly lower learning rate compared with control groups with inactive scrambled oligodeoxynucleotides (blue line) or saline (grey line) injections (Figure 1). *Post hoc* Tukey’s tests after repeated two-way ANOVA showed significantly lower success rates for the antisense group compared with scrambled oligodeoxynucleotides and saline groups on days 6 – 11 (Figure 1). In the antisense group, the fast performance improvement phase was replaced with a slow learning curve extended from day 4 to day 11 (Figure 1). However, with continued training, the success rate of the antisense group eventually reached the same asymptote as the other groups on day 12 (Figure 1). Rats with scrambled oligodeoxynucleotides or saline injection learned the skilled reaching task as efficiently as the control uninjected animals (Figure 1), indicating the surgery and intracortical injections did not affect their ability in sensorimotor learning.

**Figure 1.**
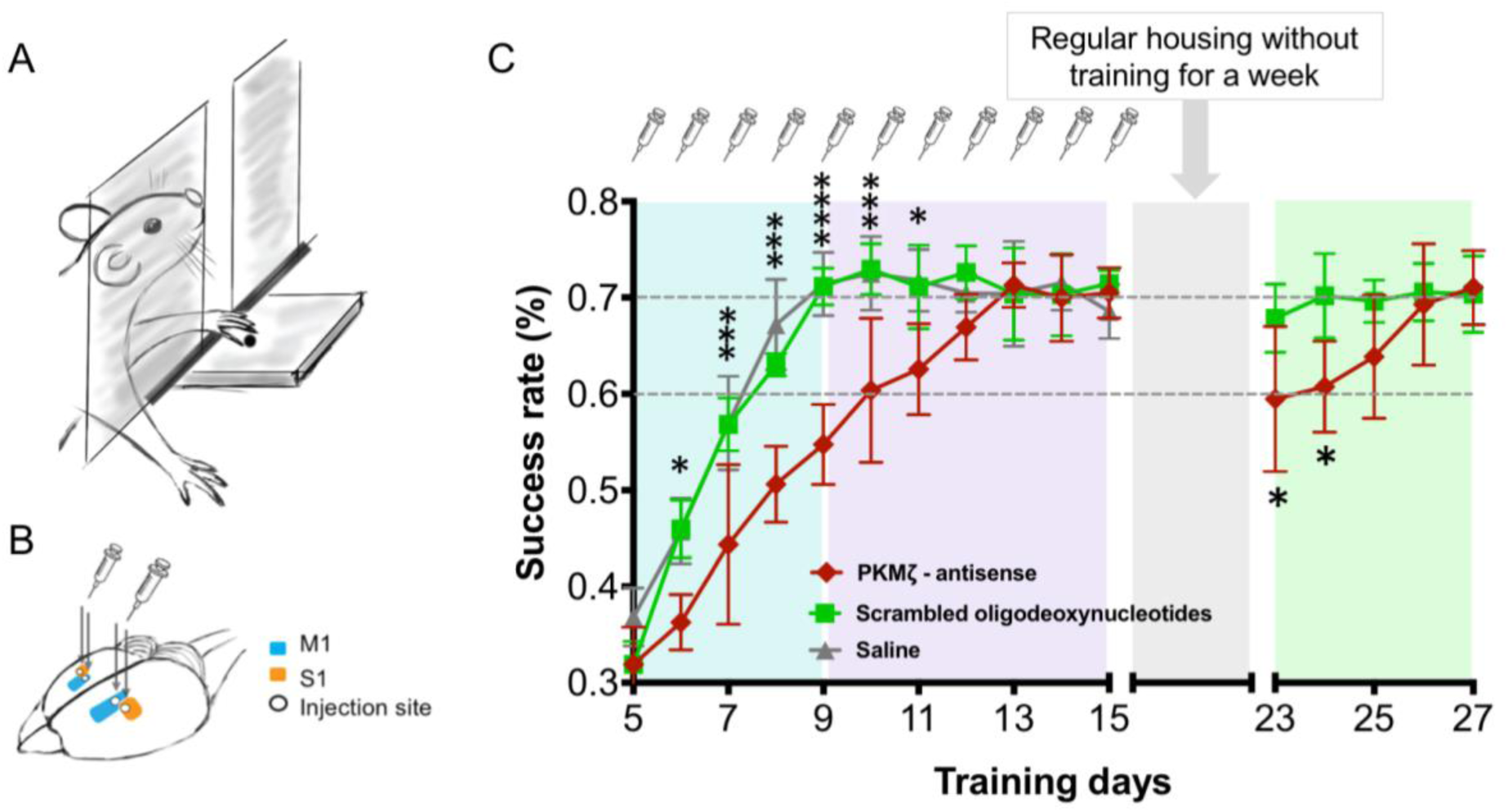
The effect of PKMζ-antisense on sensorimotor learning. **A:** Rats were trained to reach through the slot with their preferred forelimb and grasp the food pellet on the platform outside of the behavioral chamber. **B:** Bilaterally injection sites of M1 (blue) and S1(orange). **C:** PKMζ-antisense, scrambled oligodeoxynucleotides, and saline were injected bilaterally 30 minutes before the skilled reaching session every day for 15 days (1 µl/ hemisphere) in each group. The learning curves for animals that received PKMζ-antisense, scrambled oligodeoxynucleotides or saline injections were marked in red (n=6), blue (n=4) and grey (n=5) (Means ± SD). Two-way ANOVA showed significant effect between injection groups on learning (F_(2, 12)_ = 32.95; *p < 0.0001*). *Post hoc* Tukey’s tests for multiple comparisons showed significant lower success rate for the antisense group compared with scrambled oligodeoxynucleotides group and saline group on day 6 (*\*p = 0.0105, ** p = 0.0072*), day 7 (*\*\*\*p = 0.0006, ***p = 0.0002*), day 8 (*\*\*\*p = 0.0007, ****p < 0.0001*), day 9 (*\*\*\*\*p < 0.0001, **** p < 0.0001*), day 10 (*\*\*\*p = 0.0006, ***p = 0.0004*) and day 11 (*\*p = 0.0278, **p = 0.0092*). The same groups of rats that had received daily PKMζ–antisense or scrambled oligodeoxynucleotides injection in S1/M1 from day 1 to day 15, were tested again on days 23-27 after one week no training gap. Student *t-test* showed significant lower success rates of rats with PKMζ-antisense compared with scrambled oligodeoxynucleotides injections on day 23 (*\*p= 0.0313*) and day 24 (*\*p = 0.0183*). (See Figure S 3A for training from day 1-5).

### PKMζ-antisense disrupts the stability of long-term sensorimotor memory

The injection of PKMζ-antisense in S1/M1 slowed the rate of learning in the proficiency acquisition phase, but the antisense-injected animals eventually reached the same proficiency as controls with additional training days, suggesting at least two mechanisms of motor learning, one that requires *de novo* PKMζ synthesis and another that does not. Because our experiments involve continual training and reinforcement, we asked how well the latter mechanism is maintained memory without reinforcement. After asymptotic performance was achieved, we ceased training and retested the rats 1 week later. Whereas the control, scrambled oligodeoxynucleotides group maintained peak performance, as expected, the PKMζ–antisense group showed significantly lower performance on day 23 and 24 (Figure 1). We then resumed the daily training and found that the group previously injected with antisense relearned the task and reached the same asymptotic success rate again on day 26. These results indicate that new PKMζ synthesis during learning is required for the long-term maintenance of sensorimotor memory that is not reinforced by practice/reinforcement.

### ZIP specifically disrupts the storage of sensorimotor memory maintained without reinforcement

To further test this hypothesis, we examined the effect of the aPKC inhibitory peptide ZIP on sensorimotor memory maintenance after the skill was fully mastered and maintained in two schemes: one maintained with continued daily training and the other in which the memory was stored for 1 week without practice/reinforcement.

We first examined if long-term memory maintained by daily practice/reinforcement is disrupted by injections of ZIP. Rats were trained for 9 days to reach maximum proficiency in the skilled reaching task. One day after the last training session, ZIP, scrambled ZIP, or saline was injected bilaterally in S1/M1 (Figure 2A and Figure S 2). Thirty minutes after injection, reaching proficiency was tested. The success rate of rats with ZIP injection was not significantly different from those with scrambled ZIP or saline injections (Figure 2A). The training was continued for the next two days and no further difference was found. To confirm that memories held by daily practice and reinforcement were not affected by ZIP, a second injection was given on day 13 in the same rats, and 1 day later the training resumed from days 14-16. Again, ZIP had no effect on the task performance (Figure 2A).

**Figure 2.**
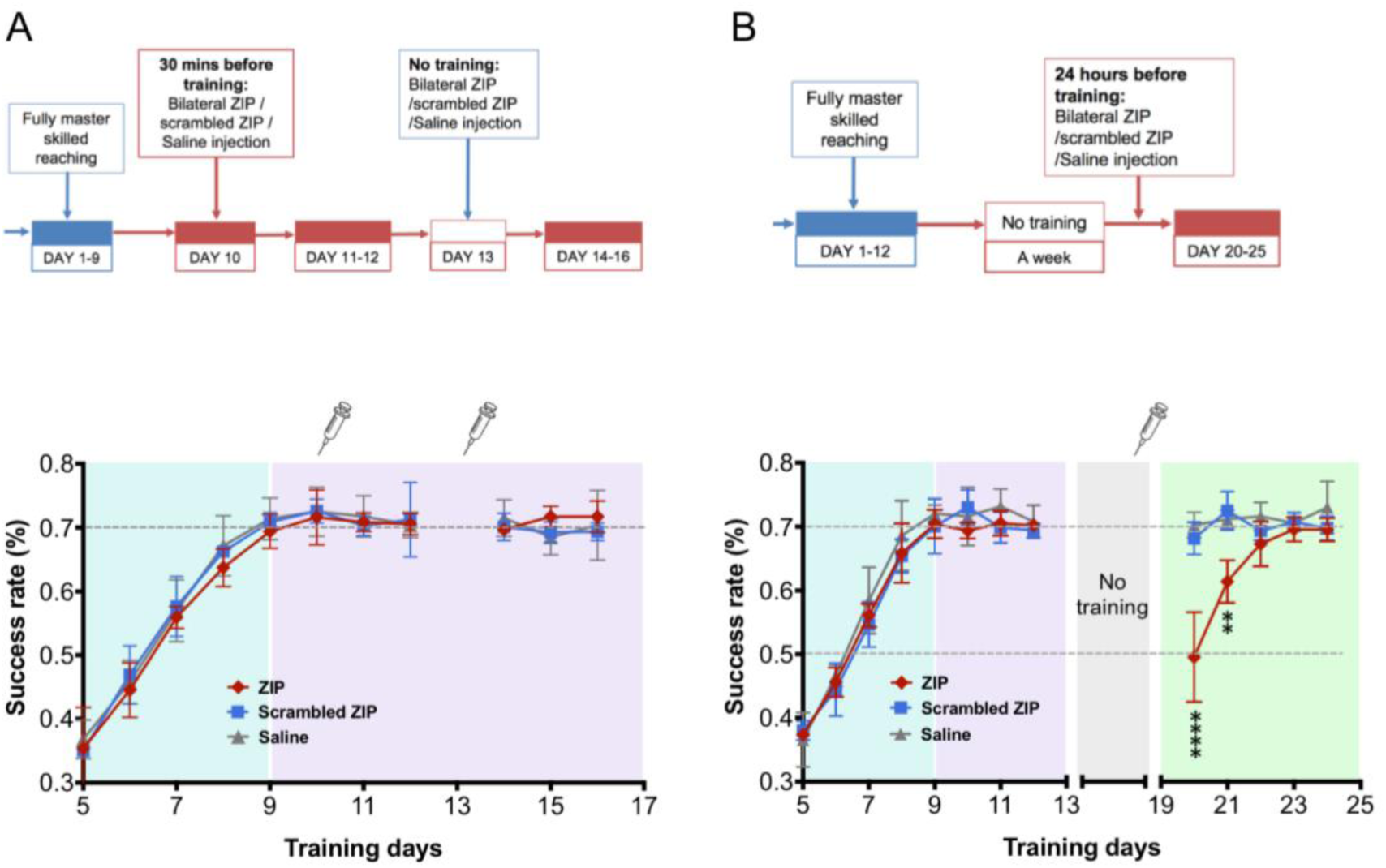
The effect of ZIP on motor memory maintenance. **A:** Motor memory sustained by continual practice is PKMζ-independent. Rats were trained for the skilled reaching for 9 days until the success rate reached plateau: ~70%. On day 10, the well-trained animals were randomly divided into 3 groups to receive ZIP/scrambled ZIP/ saline injection bilaterally 30 minutes before the skilled reaching task. The injection dosage of ZIP/scrambled ZIP was 1µl per site and 2 µl in total on each hemisphere. Daily training was continued for day 11 and 12. On day 13, the same injections were made to the same rats as day 10, but without training. Daily training was resumed from day 14-16 and the behaviors were recorded as the successful reaches divided by the total reaches in each session. The learning curves for rats that received ZIP, scrambled ZIP or saline injections were marked in red (n = 3), blue (n=3) and grey (n = 5) respectively (Means ± SD). Two-way ANOVA showed no significant difference of learning rate between groups (F_(2,8)_ = 0.6555, *p = 0.5450*). (See Figure S 3B for training from day 1-5). **B:** Motor memory sustained without practice is PKMζ-dependent. Rats were trained daily for 12 days followed by one week regular housing, and received ZIP, scrambled ZIP, or saline injections bilaterally on day 19 (2 µl / hemisphere). Since day 20, daily training was resumed and the behaviors were recorded as the successful reaches divided by the total reaches in each session. The learning curves for rats that received ZIP, scrambled ZIP and saline injections were marked in red (n=5), blue (n=5) and grey respectively (Means ± SD). Two-way ANOVA (F(2,8) = 6.927, p = 0.0180), followed by post hoc tukey’s comparisons showed significant lower success rate of ZIP on day 20 and 21 compared with scrambled ZIP (\*\*\*\**p*<0.0001 and \*\**p*=0.0024), and saline groups (\*\*\*\**p*<0.0001 and \*\**p*=0.0086). (See Figure S 3C for training from day 1-5).

A second set of rats were trained to asymptotic performance and after 1 week of no training were divided into 3 groups that received either ZIP, scrambled ZIP, or saline injection bilaterally in S1/M1 (Figure 2B and Figure S 2). Rats intracortically injected with scrambled ZIP or saline showed no loss of proficiency in the reaching task when tested again on day 20 (1 day after injection). In striking contrast to the reinforced memory, ZIP injections disrupted long-term motor memory that was maintained without practice/reinforcement (Figure 2B). On resuming daily training from day 20-25, a relearning curve revealed that group previously injected with ZIP acquired motor memory with proficiency indistinguishable from their original learning curve, indicating no saving of motor memory after memory erasure by ZIP.

### Sensorimotor training induces a persistent increase of PKMζ in sensorimotor cortex maintained during long-term memory storage

To localize the persistent increase in PKMζ that maintains long-term motor memory, we next measured throughout S1 and M1 forelimb regions during each learning phase. Confocal microscopy revealed PKMζ in the sensorimotor cortex is compartmentalized in small puncta (Figure 3), similar to its distribution in hippocampus [36]. The puncta number and size were quantified in each hemisphere and the interhemispheric ratios (trained / untrained hemispheres) were used to compare the levels of PKMζ in each group with controls (naïve rats) (Figure 4 and Figure S 4).

**Figure 3.**
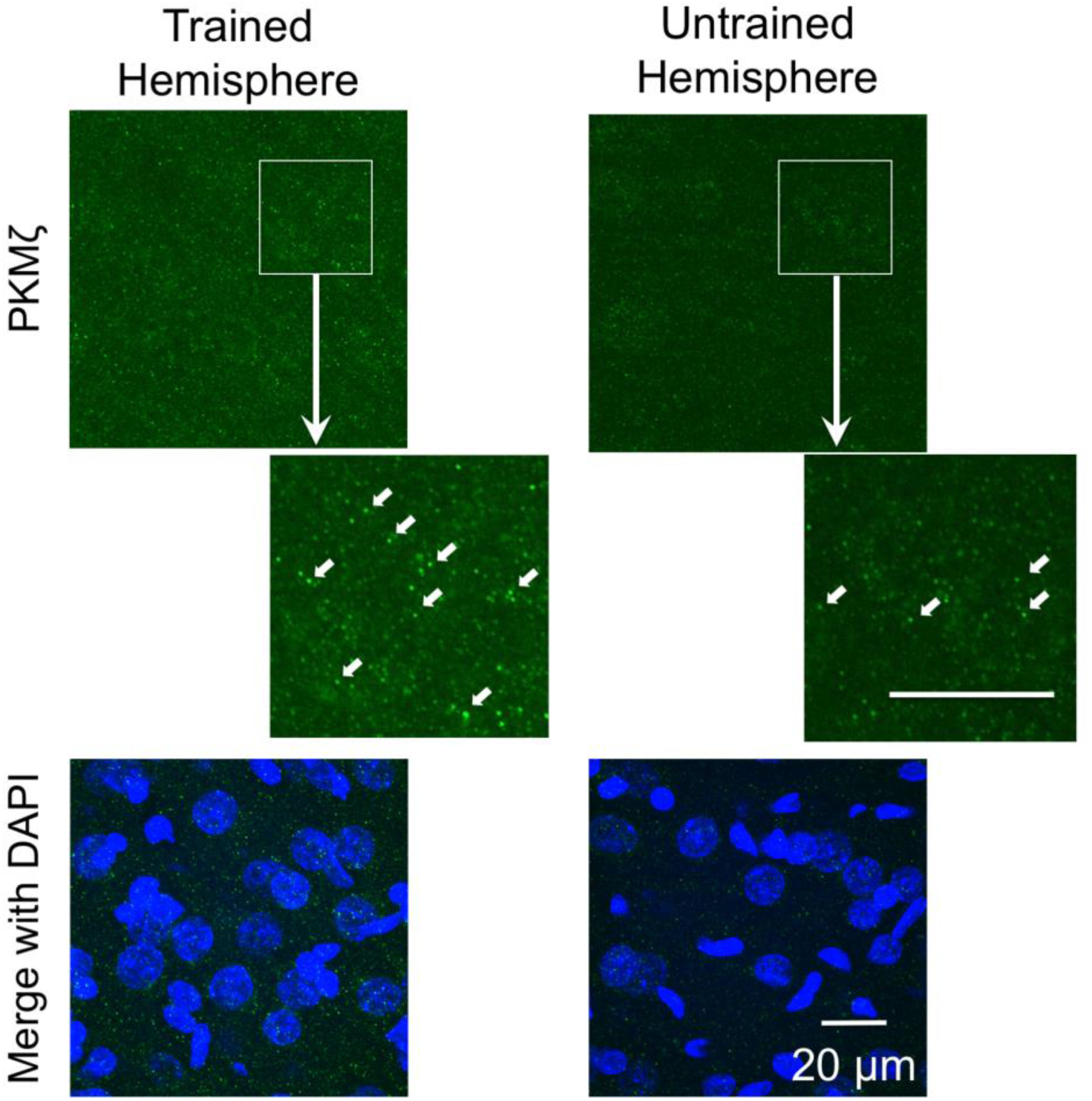
PKMζ immunostaining of sensorimotor cortex. Immunostaining of PKMζ could be detected as small puncta (indicated by the white arrows). Green - PKMζ staining; Blue – DAPI staining; Scale bar, 20 μm. Example images were from M1 layer III/III of a rat with 9 days of training (magnification 120x).

**Figure 4.**
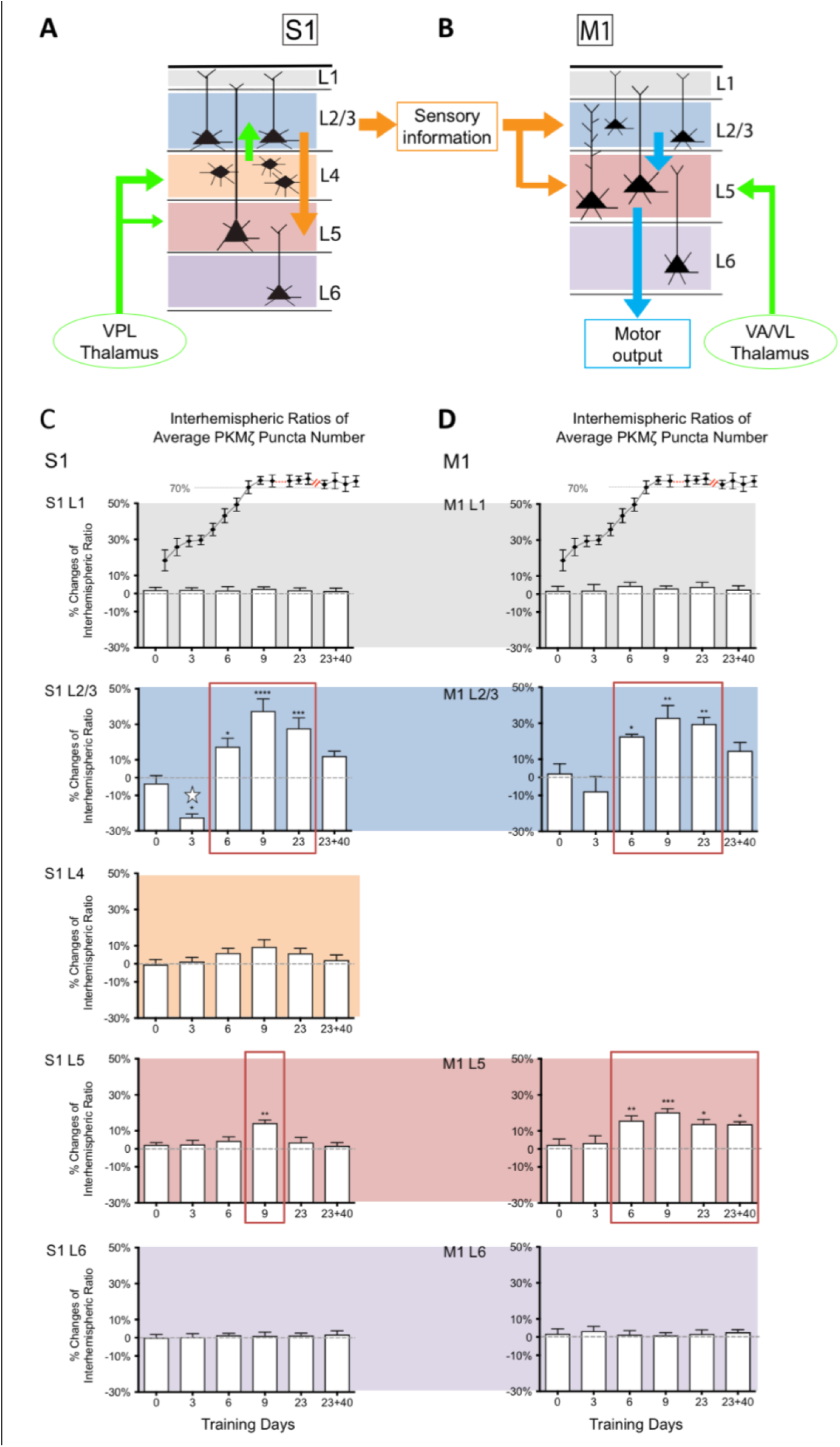
Spatiotemporal Changes of PKMζ Puncta Number in Sensorimotor Learning. **A, B**: Information propagation in S1 and M1 during sensorimotor learning. **C, D**: Layer-specific changes of PKMζ puncta number ratios in S1 and M1. X-axis - days of training; Y-axis – the % changes of interhemispheric ratio of average PKMζ puncta number (Means ± SEM). One-way ANOVA showed significant changes in S1 layer II/III (F(_5,26_) = 20.55; *p*<0.0001), S1 layer V (F(_5,26_) = 5.079; *p*=0.0022), M1 layer II/III (F(_5,26_) = 8.764; *p*<0.0001) and M1 layer V (F(_5,26_) = 6.857; *p*=0.0003). Dunnett’s multiple comparison test showed the difference of each training group with control (* p<0.05, ** p<0.01, ***p<0.001, ****p<0.0001). In contrast to puncta number, the interhemispheric ratios of the average PKMζ puncta size did not change significantly (see Figure S 4).

The results revealed that learning-induced changes in PKMζ, as well as the persistence of these changes during memory maintenance, are selective to distinct cortical layers. One-way ANOVA showed significant changes of PKMζ puncta numbers in both S1 and M1 layers II/III and V. In the initial skill acquisition phase, which is not affected by PKMζ-antisense or ZIP, the interhemispheric ratios of PKMζ puncta number decreased significantly compared with control in S1 layer II/III (Figure 4C). In contrast, during the performance improvement that is disrupted by these PKMζ antagonists, PKMζ increased in multiple cortical layers. After 6 and 9 days of training, PKMζ puncta numbers increased in S1 layer II/III, M1 layer II/III and M1 layer V, and, after 9 days of training, in S1 layer V (Figure 4C, D). The increases reached a maximum on day 9 when memory expression reached asymptotic levels of performance. During the proficiency maintenance phase, when the animals continued daily training, the increased amounts of PKMζ in S1 layer II/III, M1 layer II/III and M1 layer V were maintained, while that in S1 layer V returned to basal levels (Figure 4C, D).

During the long-term memory storage phase, i.e., after 40-days without training, the increases in PKMζ persisted specifically in M1 layer V (Figure 4C, D). As a control, we examined in all S1/M1 cortical layers a second protein associated with learning-induced structural changes in sensorimotor cortex, postsynaptic density protein-95 (PSD-95). In contrast to PKMζ, PSD-95 increased transiently only at the end of performance improvement training phase and did not persistent beyond that phase (Figure S 5). These results demonstrate that skilled motor training induces selective increases in PKMζ that at long time scales can be either transient or highly stable within specific layers of sensorimotor cortex.

## Discussion

Here we show for the first time that persistent increases in PKMζ store long-term skilled motor memory that is maintained without reinforcement, and the persistence of this molecular mechanism of long-term memory is found not throughout sensorimotor cortex but specifically in the output cortical layer of M1.

Both PKMζ-specific antisense oligodeoxynucleotides and the aPKC-selective kinase inhibitor disrupted the maintenance of long-term skilled motor memory for the reaching task that is maintained without practice, but not skilled motor memory that is reinforced daily. These results are in line with earlier findings with ZIP on conditioned taste aversion (CTA), in which the inhibitor erased stored CTA memory, but had no effect on CTA memories that were recently reconsolidated by re-exposure to the conditioning stimulus [37]. However, ZIP cannot exclude the possibility of an additional role for PKCζ [33]. Our results indicate that persistent practice/reinforcement and another mechanism of memory can maintain the expression of enhanced motor performance for ~1 day. Such mechanisms might include persistent action of other PKC isoforms or other kinases such as CaMKII.

The changes of PKMζ in the forelimb contralateral S1 layer II/III are biphasic. During early skill acquisition, PKMζ is initially downregulated in S1 layer II/III (Figure 4C), which may represent a weakening of the sensory map or sensory-motor associations during the initial phase of sensorimotor learning, when there is a pause in performance improvement prior to rapid acquisition of the stable motor engram (Figure S 1) [8,32,38]. Downregulation of PKMζ is associated with long-term depression [39,40], and, therefore, the initial decrease of PKMζ in S1 layer II/III might represent an LTD-like process involving a weakening of pre-existing neuronal networks. Because rats were still actively exploring the training chamber and food pellet, each reaching attempt at this phase could induce new sensory stimuli patterns and the comparatively low success rate might act as negative feedback to disrupt previously acquired sensory-motor associations. This initial phase of motor learning is unaffected by PKMζ antagonists.

In contrast, during the performance improvement phase that begins after the 5^th^ day of training, the amount of PKMζ rebounds and increases above baseline in S1/M1 layers II/III and V (Figure 4C, D and Figure S 4). The timing of the increase of PKMζ parallels the LTP-like potentiation of synaptic transmission observed in motor cortex after skill learning [8,9]. PKMζ -antisense specifically delays the acquisition of the sensorimotor memory phase, in line with PKMζ’s critical role in maintaining LTP.

The persistence of the learning-induced PKMζ increases are layer-specific, revealing the location of the very long-term molecular mechanism maintaining motor memory. After 40 days without reinforcement, the PKMζ increases in sensory cortex and layers 2/3 of motor cortex return towards baseline. In contrast, the long-term increases in PKMζ in layer V of motor cortex are highly stable, indicating that the persistent molecular changes associated with stable skilled motor memory are within the cellular output circuitry of primary motor cortex. This localization is in line with work showing changes to thalamocortical synaptic plasticity that target cortical layer V neurons projecting to C8 and the distal forepaw muscles used in learned grasping on this rodent reaching task [41]. Further research will be required to determine the particular synapses within cortical layer V that express increased amounts of PKMζ to maintain procedural memories for long periods of time.

## Author Contributions

P.P.G. performed the experiments and analyzed the data. J.T.F., J.H.G. and T.C.S. supervised the project and provided input on experimental design. All the authors listed wrote the manuscript together.

## Acknowledgments

We would like to thank Dr. Changchi Hsieh, Dr. Janina Ferbinteanu and Dr. Panayiotis Tsokas for technical advice, and thank all the members of the Francis lab for discussion and support. This study was supported by grants from DAPAR (www.darpa.mil) (Award#60806; Project#:108723) (J.T.F.) and IBR-SUNY Downstate graduate fund (J.T.F and J.H.G.).

## Supplemental

**Figure S1** (refers to Results: Learning phases of a skilled reaching task)

**Figure S 1.**
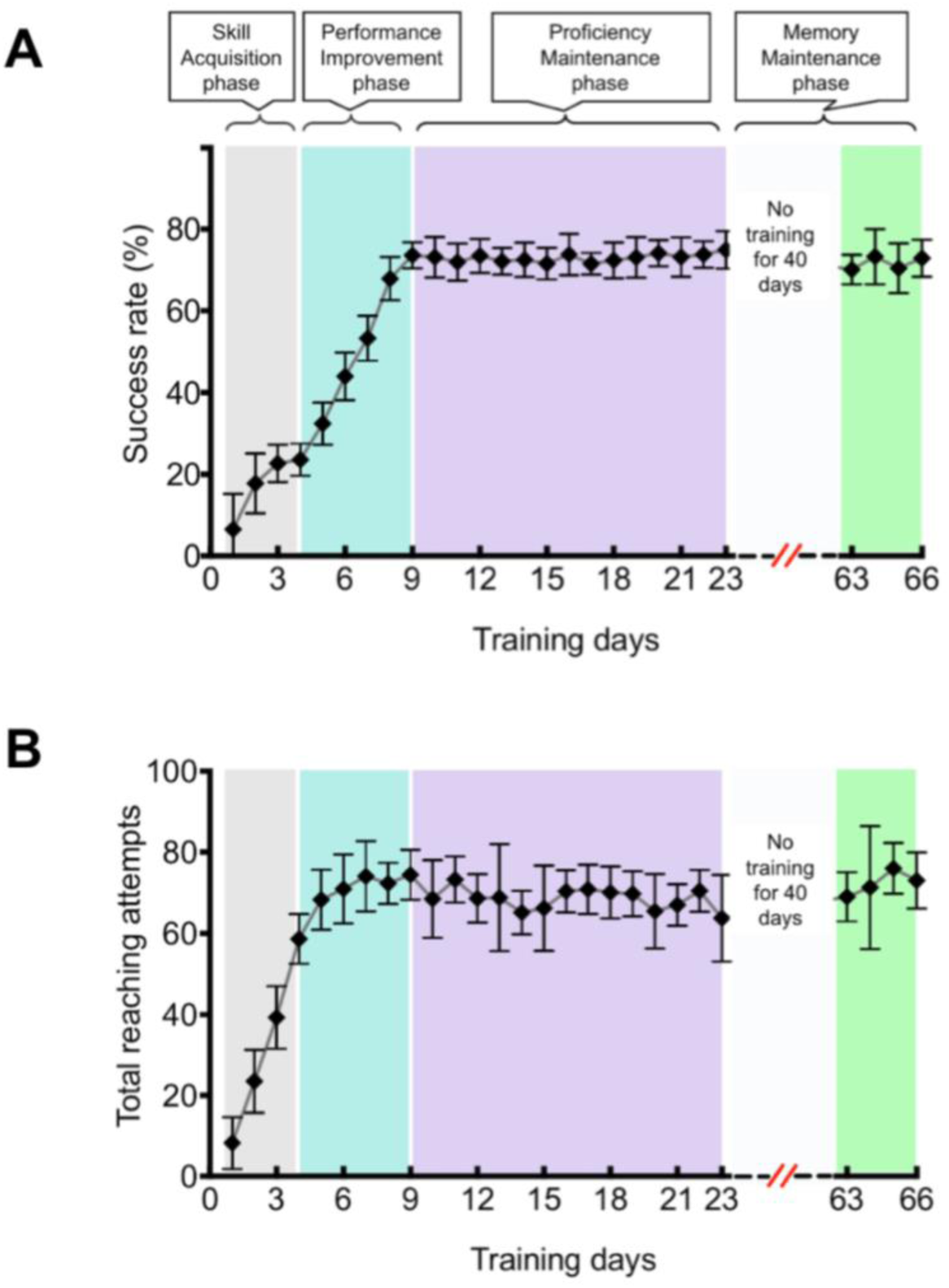
Daily success rate and total reaching attempts in the skilled reaching task. **A:** The learning curve for skilled reaching task. X-axis – the number of training days; Y-axis – success rate (%): successful reaches divided by the total reaching attempts (Means ± SEM). **B:** The total reaching attempts in a daily 30 minutes training session. X-axis – the number of training days; Y-axis – total reaching attempts: the average number of total reaching attempts (Means ± SEM); Grey block - the skill acquisition phase; Blue block - the performance improvement phase; Purple block - the proficiency maintenance phase; White block - regular housing without training for 40 days; Green block - testing after 40 days no training gap, and was defined as the memory maintenance phase together with white block.

**Figure S2** (refers to Results PKMζ-antisense slows the performance improvement phase of sensorimotor learning and ZIP specifically disrupts the storage of sensorimotor memory maintained without reinforcement)

**Figure S 2.**
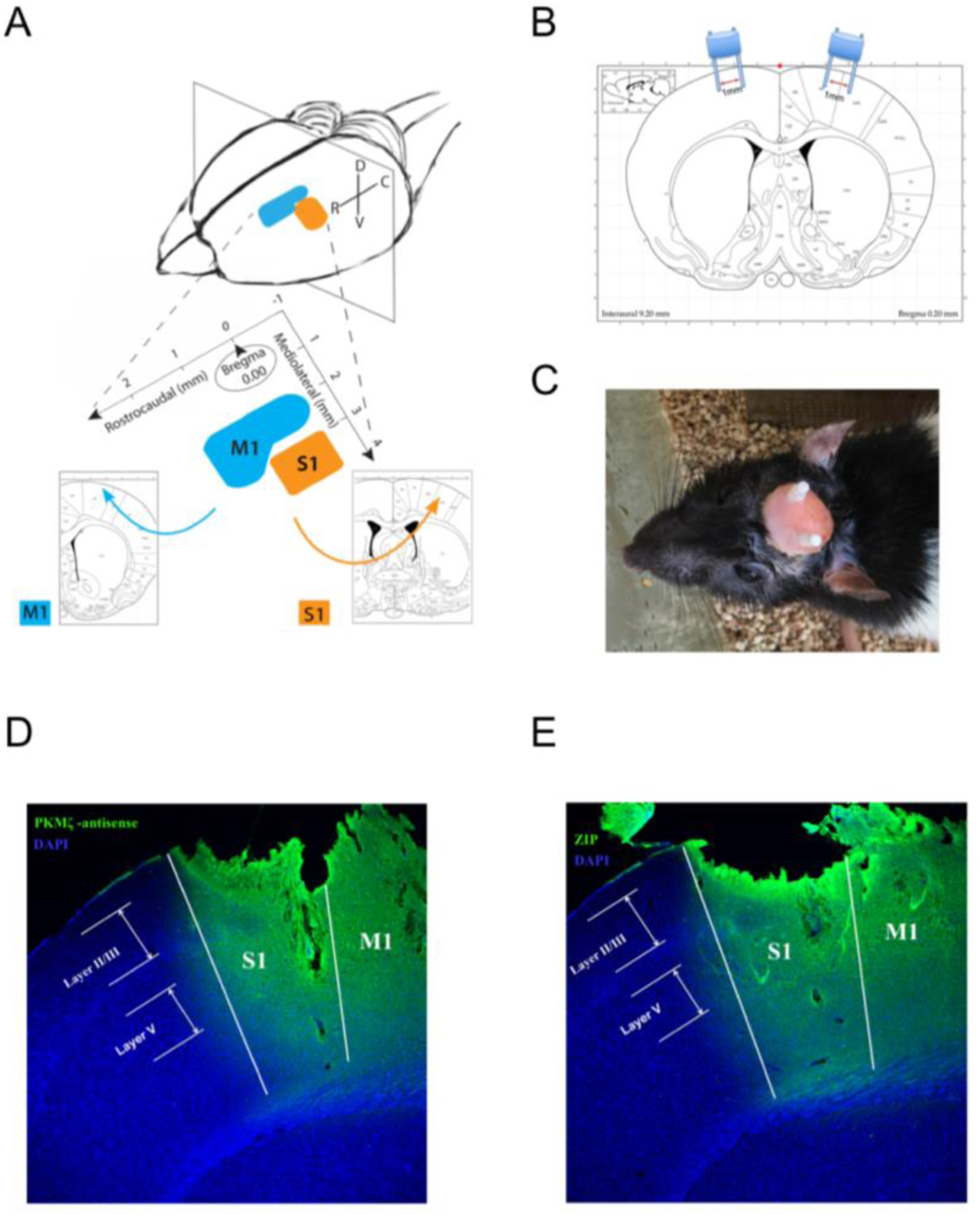
Fluorescence labeling of biotinylated PKMζ-antisense injection. **A:** The stereotaxic coordinates for rat forelimb M1 and S1 regions. The coronal sections for IHC study were chosen from primary motor cortex (M1, 1.6 mm ~ −0.5 mm to bregma; 1.5 mm ~ 3 mm lateral to the midline) and primary somatosensory cortex (S1, 0.5 mm ~ −0.4 mm to bregma; 3 mm ~ 4mm lateral to the midline) in rat. **B:** Demonstration of the location for implanted double guide cannula bilaterally. The center-to-center distance between two dummy wires in each double guide cannula was 1.0 mm. They were tilted 10˚ on each hemisphere from the vertical line. The coordination of guide cannula in M1: AP + 0.2 mm, ±ML 2.2 mm, DV −1.16 mm; The coordination of guide cannula in S1: AP + 0.2 mm, ±ML 3.2 mm DV - 0.99 mm. **C:** Top view of a rat 12 hours after surgery with cannula implantation. **D:** Fluorescence labeling of biotinylated PKMζ-antisense injection. Rat was sacrificed 1 hour after the injections of biotinylated PKMζ-antisense in PBS. The injection dosage was 0.5μl/site and 1 μl per hemisphere in total. DAPI was counterstain. It was used to evaluate the spread of PKMζ-antisense and prove the effectivity of our injection approach. Green: biotinylated PKMζ-antisense. Blue: DAPI. E: Fluorescence labeling of biotinylated ZIP injection. Rat was sacrificed 1 hour after the injections biotinylated ZIP solution. The injection dosage was 1 μl/site and 2 μl per hemisphere in total. DAPI was counterstain. It was used to evaluate the spread of ZIP and prove the effectivity of our injection approach. Green: biotinylated ZIP. Blue: DAPI.

**Figure S3** (refers to Results PKMζ-antisense slows the performance improvement phase of sensorimotor learning and ZIP specifically disrupts the storage of sensorimotor memory maintained without reinforcement)

**Figure S 3.**
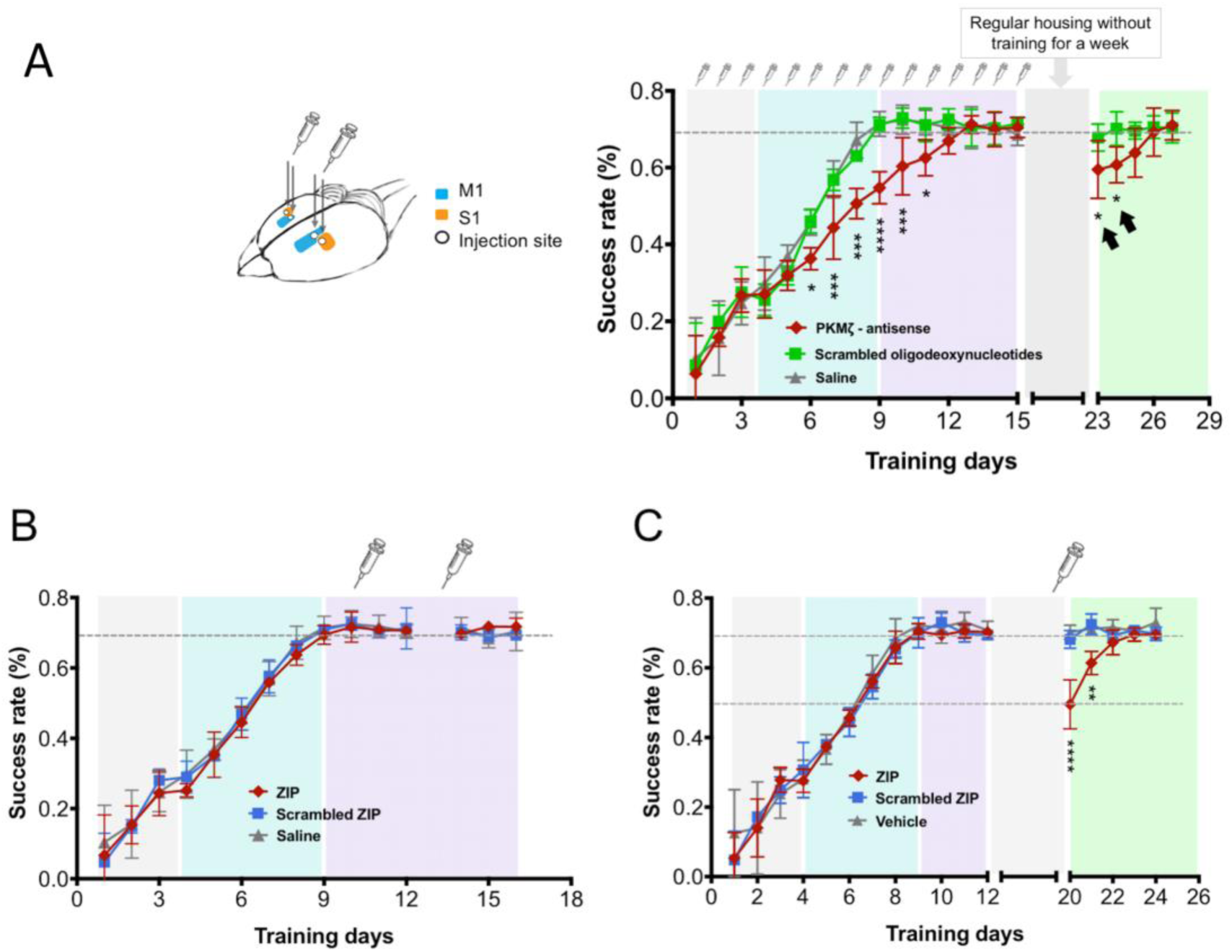
Perturbing PKMζ synthesis and aPKC activity during motor learning and memory maintenance. A: the complete learning curve for Figure 1. The effect of PKMζ-antisense on sensorimotor learning. B, C: the complete learning curve for Figure 2. The effect of ZIP on motor memory maintenance.

**Figure S4** (refers to Results: Sensorimotor training induces an initial decrease followed by a persistent increase of PKMζ in sensorimotor cortex that is maintained during long-term memory storage)

**Figure S 4.**
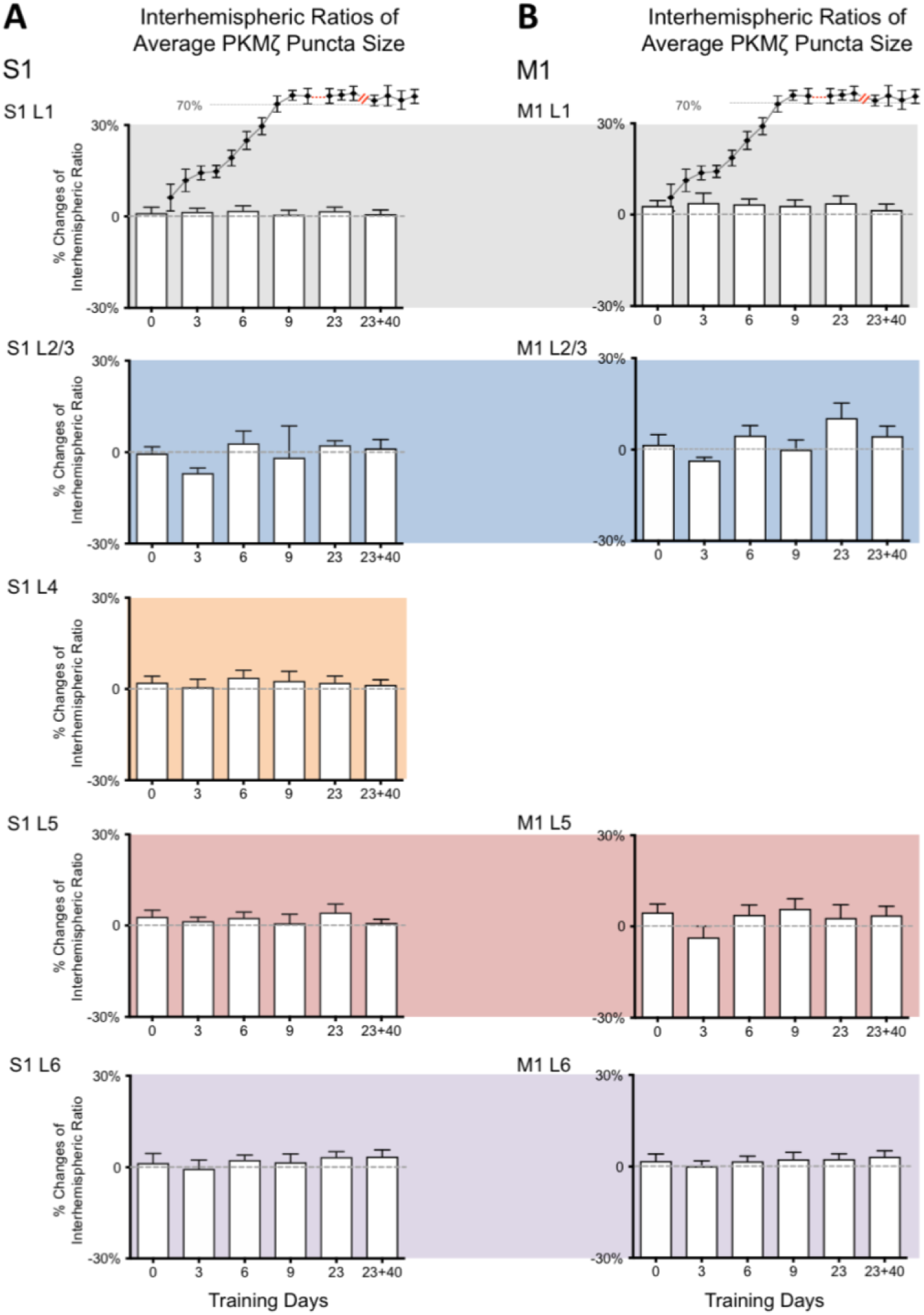
Spatiotemporal Changes of PKMζ Puncta Size in Sensorimotor Learning. **A, B:** The layer-specific changes of average PKMζ puncta size in S1 and M1 along with motor learning. X-axis - days of training (0 - naïve rat; 3, 6, 9, 23 - rats trained for 3, 6, 9 or 23 days; 23+40 – rats trained for 23 days and regular housed for another 40 days); Y-axis – the % of interhemispheric ratio of averaged PKMζ puncta number (Means ± SEM); No significant between group effects were found with one-way ANOVA in all the regions of both S1 and M1.

**Figure S5** (refers to Results: Sensorimotor training induces a persistent increase of PKMζ in sensorimotor cortex maintained during long-term memory storage)

**Figure S 5:**
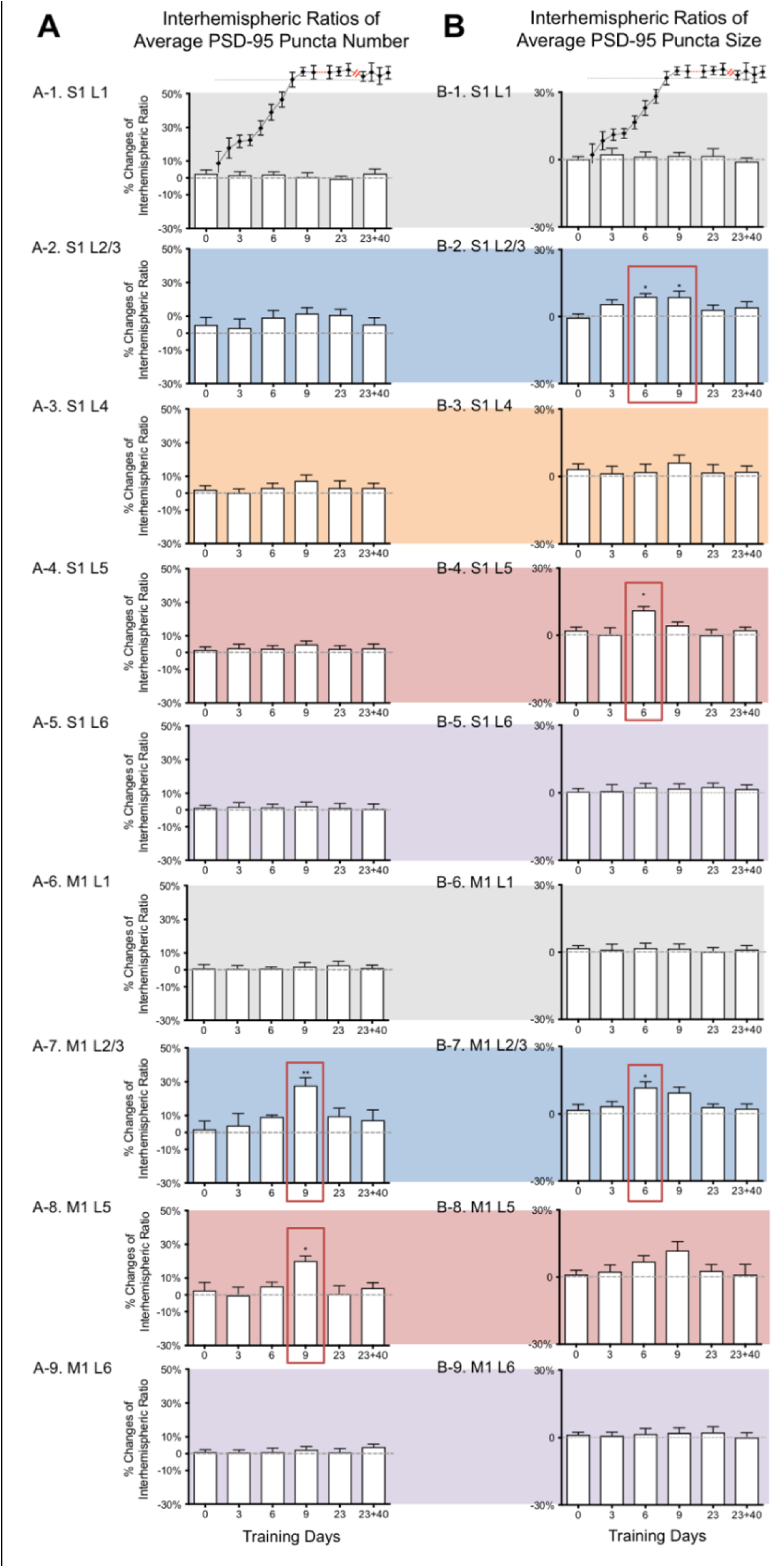
Spatiotemporal Changes of PSD-95 Puncta Number and Size in Motor Learning. **A**: The layer-specific changes of PSD-95 puncta number in S1 and M1 along with motor learning. **B**: The layer-specific changes of PSD-95 puncta size in S1 and M1 along with motor learning. A-1 and B-1: S1 layer I; A-2 and B-2: S1 layer II/III; A-3 and B-3: S1 layer IV; A-4 and B-4: S1 layer V; A-5 and B-5: S1 layer VI; A-6 and B-6: M1 layer I; A-7 and B-7: M1 layer II/III; A-8 and B-8: M1 layer V; A-9 and B-9: M1 layer VI. X-axis - the days of training (0 - naïve rat; 3, 6, 9, 23 - rats trained for 3, 6, 9 or 23 days; 23+40 – rats trained for 23 days and regular housed for another 40 days); Y-axis - the % of interhemispheric ratio of average PSD-95 puncta number (A) and average PSD-95 puncta size (B) (Means ± SEM); Statistical analysis was conducted using one-way ANOVA for between group effects, followed by Dunnett’s multiple comparison test to show the difference of each training group with control (0 - naïve rat) (* *p*<0.05, ** *p*<0.01, \*\*\**p*<0.001, \*\*\*\**p*<0.0001).

